# Dating ancient human samples using the recombination clock

**DOI:** 10.1101/023341

**Authors:** Priya Moorjani, Sriram Sankararaman, Qiaomei Fu, Molly Przeworski, Nick Patterson, David Reich

## Abstract

The study of human evolution has been revolutionized by inferences from ancient DNA analyses. Key to these is the reliable estimation of the age of ancient specimens. The current best practice is radiocarbon dating, which relies on characterizing the decay of radioactive carbon isotope (^14^C), and is applicable for dating up to 50,000-year-old samples. Here, we introduce a new genetic method that uses recombination clock for dating. The key idea is that an ancient genome has evolved less than the genomes of extant individuals. Thus, given a molecular clock provided by the steady accumulation of recombination events, one can infer the age of the ancient genome based on the number of missing years of evolution. To implement this idea, we take advantage of the shared history of Neanderthal gene flow into non-Africans that occurred around 50,000 years ago. Using the Neanderthal ancestry decay patterns, we estimate the Neanderthal admixture time for both ancient and extant samples. The difference in these admixture dates then provides an estimate of the age of the ancient genome. We show that our method provides reliable results in simulations. We apply our method to date five ancient Eurasian genomes with radiocarbon dates ranging between 12,000 to 45,000 years and recover consistent age estimates. Our method provides a complementary approach for dating ancient human samples and is applicable to ancient non-African genomes with Neanderthal ancestry. Extensions of this methodology that use older shared events may be able to date ancient genomes that fall beyond the radiocarbon frontier.

**Significance:** We introduce a new genetic method for dating ancient human samples that uses the recombination clock. The main idea relies on the insight that an ancient genome lacks several thousand years of evolution compared to genomes of living individuals. To infer the age of ancient genomes, we take advantage of the shared history of Neanderthal gene flow into non-Africans that occurred around 50,000 years ago. By characterizing the dates of Neanderthal gene flow in ancient and extant genomes and quantifying the difference in these dates, we estimate the age of the ancient specimen. Our method is applicable for dating ancient samples more recent than the Neanderthal mixture event, so on par with radiocarbon dating, providing a complementary approach for dating.

## Introduction

Ancient DNA analyses have transformed research into human evolutionary history, providing an unprecedented opportunity to learn about genetic patterns present in the past (1). In order to properly interpret findings from an ancient sample, it is important to have an accurate estimate of the age of the specimen. The standard tool used for this purpose is *radiocarbon dating*, which relies on measuring the decay of radioactive carbon isotope ^14^C over time, and is applicable for dating samples up to 50,000 years old (2). This method is based on the principle that when a living organism dies, the existing amount of ^14^C starts decaying at a constant rate, with a half-life of 5,730 ± 40 years (3, 4). By measuring the ratio of ^14^C to ^12^C (stable form) in the sample and assuming that the starting ratio of carbon isotopes is the same everywhere in the biosphere, one can infer the age of the sample. A complication is that carbon isotope ratios vary between reservoirs (e.g. marine, freshwater, atmosphere) and over time (2, 5). Thus, radiocarbon dates must be converted to calendar years using calibrations curves, which are constructed by dating materials with ^14^C and alternate methods such as annual tree rings (dendrochronology) or Uranium-series dating of coral (2). Such calibrations, however, may not fully capture the variation in atmospheric carbon at historical timescales. In addition, contamination of a sample by modern carbon, introduced during excavations or while dating, can bias the inferred dates (2). Other methods such as stratigraphy, bone nitrogen and potassium–argon dating also exist; however these suffer from similar drawbacks, relying on calibration data that are often difficult to obtain (6).

Here, we describe an alternative approach for dating ancient samples using genetic data, applicable in cases where DNA sequence data are available, as is becoming increasingly common (1). This method relies on the insight that an ancient genome has experienced fewer generations of evolution compared to the genomes of its living relatives. Since recombination occurs at a constant rate per generation, the accumulated number of recombination events provides a molecular clock for the time elapsed or in the case of an ancient sample, the number of missing generations since it ceased to evolve (7). This idea is referred to as “branch shortening” and estimates based on branch shortening can be translated into absolute time in years by using an independent calibration point such as divergence time in years as well as an estimate of the average human generation interval, i.e., the mean age of reproduction.

Branch shortening has previously been used in wide-range of applications in genetics for dating phylogenies, inferring mutation rates, and improving inferences of population history in humans and other species (8-10). The idea of using branch shortening to date ancient samples was first applied on a genome-wide scale by Meyer et al. 2012, who used the mutation clock (instead of the recombination clock as proposed here) to estimate the date of the Siberian Denisova finger bone, which is probably older than 50,000 years, and has not been successfully radiocarbon dated (11). Specifically, these authors compared the sequence divergence between the Denisova and present-day human genomes, and calibrated the branch shortening relative to human-chimpanzee divergence time (11). The use of ape divergence times as calibration point, however, relies on estimates of mutation rate that are uncertain. In particular, recent pedigree-based sequencing studies have yielded a mutation rate per year that is approximately two fold lower than the one obtained from phylogenetic methods (12, 13). In addition, comparison to human-chimpanzee divergence time relies on branch shortening estimates that are very small relative to the total divergence of millions of years, so that small errors in mutation calling can bias estimates. This introduces substantial uncertainty in the estimates of the age of the ancient samples, making this approach impractical for dating samples that are only tens of thousands of years old, a time period that encompasses the vast majority of ancient human samples sequenced to date.

Given the challenges associated with the use of the mutation clock, here we explore the possibility of using the molecular clock based on the accumulation of crossover events (recombination clock). The crossover landscape in humans is well characterized and current genetic maps have high accuracy even at short distances (e.g. 10-50 Kb) (14). In addition, instead of using a distant outgroup such as chimpanzees, we rely on a more recent shared event that has affected both extant and ancient modern humans and hence a more reliable fixed point on which to base the dating. As the vast majority of ancient samples sequenced to date were discovered in Eurasia (with estimated ages: ∼2,000-45,000 years before present (yr BP)), post-date the Neanderthal admixture, and show evidence of admixture with Neanderthals between 37,000–86,000 yr BP, we use the Neanderthal gene flow as the shared event (15, 16).

To estimate the age of the ancient genome, we first estimate the dates of Neanderthal gene flow in ancient and extant genomes. Because the ancient sample is closer to the shared Neanderthal admixture event (due to branch shortening), we should obtain more recent dates of Neanderthal admixture in ancient samples compared to extant samples. The difference in the inferred dates is thus informative about the age of the ancient genome. An illustrative model is shown in Figure 1. We note that an implicit assumption here is that the Neanderthal admixture into the ancestors of modern human occurred approximately at the same time, and the same interbreeding events contributed to the ancestry of all the non-African samples being compared. This method is thus not applicable for dating genomes that do have a history of Neanderthal ancestry, such as sub-Saharan African genomes.

**Figure 1:**
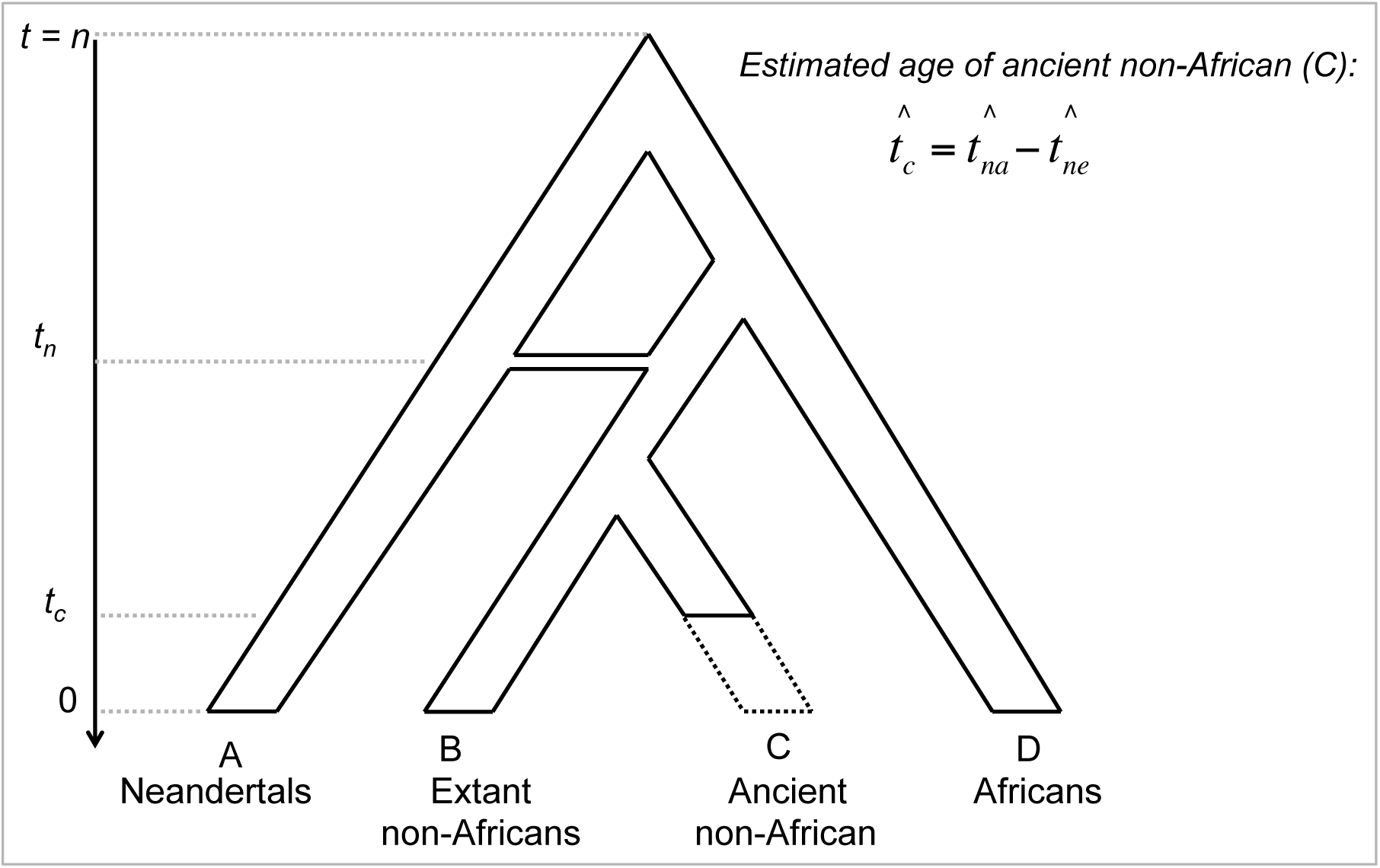
The model underlying our inference of the age of ancient genomes. We assume a simple demographic history relating Neanderthals, non-Africans and Africans. Neanderthal gene flow into non-African ancestors occurred *t*_*n*_ generations ago. This event was shared among all non-Africans and did not affect Africans. The ancient non-African genome was sampled at time *t*_*c*_. To estimate the age of the ancient genome, we first estimate the dates of Neanderthal gene flow in ancient genomes 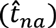 and extant genomes 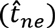. The difference in the inferred dates provides an estimate of the age of the ancient sample 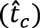.

To date the Neanderthal gene flow, we use the insight that gene flow between genetically distinct populations, such as Neanderthals and modern humans, creates correlation in ancestry across the genome that breaks downs at a constant rate per generation as crossovers occur (17-19). Thus, by jointly modeling the decay of Neanderthal ancestry and recombination rates across the genome, we can estimate the date of Neanderthal gene flow (17). Similar ideas have been previously used to date admixture events between contemporary human populations (18-20) as well as between Eurasians and Neanderthals (16, 21). Using an estimate of the human generation time, we then convert the branch-shortening estimate in generations into the age of the ancient genome in years. This method is applicable for dating ancient samples that are more recent than the Neanderthal admixture and thus applicable over the same time range as ^14^C dating.

An important feature of our method is that it is expected to give more precise results for samples that are older, as the older samples are closer to the date of Neanderthal introgression which we use for calibration, making it easier to accurately date the Neanderthal admixture. Also the magnitude of difference in Neanderthal dates between the extant and ancient samples increases as the sample becomes older, and therefore the estimated age is expected to become more precise for older samples. Thus, unlike ^14^C dating, the genetic approach becomes more reliable with age and in that regard complements ^14^C dating.

## Results

### Model and simulations

To estimate the age of ancient genomes that have Neanderthal ancestry, we developed a new method that estimates the date of Neanderthal gene flow in ancient and extant genomes first and then quantifies the difference in dates to infer the age of the ancient genome. Most current methods require multiple genomes sampled from the target population and are not applicable for dating admixture in a single diploid genome (as required here for dating ancient genomes). Thus, we took advantage of our recently developed method described in Fu et al. 2014 that computes the ancestry covariance across sites in a single diploid genome to date the Neanderthal introgression in ancient genomes (rather than the admixture related allelic correlation i.e., admixture linkage disequilibrium (LD) used by the population-based methods) (21). Specifically, this method measures the extent of covariance across pairs of markers of putative Neanderthal ancestry, i.e. sites where Neanderthals carry at least one derived allele (relative to the chimpanzees) and all individuals in a panel of sub-Saharan Africans (which have little or no evidence of Neanderthal ancestry (22)) carry the ancestral allele (21). We choose this ascertainment because it minimizes the signal of background correlation, while amplifying the signal of Neanderthal ancestry (16). This statistic (referred to as the single-sample statistic henceforth) is expected to decay approximately exponentially with genetic distance and the rate of decay should be informative of the time of gene flow (21). Assuming the gene flow occurred as a single event and fitting a single exponential to the decay pattern, we estimated the date of the Neanderthal gene flow in the target genome. Violations of this assumption can lead to inaccurate dates, and we revisit this later with examples from real data.

To test the accuracy and precision of our Neanderthal dating strategy, we performed coalescent simulations by constructing genomes of Neanderthals, west Africans and Eurasians, with Eurasians deriving 3% of their ancestry from Neanderthal gene flow (16) and the time of gene flow set to values between 100-2,500 generations ago which includes most of the time-depths relevant to our application. Parameters of the simulations were chosen such that population differentiation (F_ST_) between west Africans and Eurasians and the D-statistic between west Africans, Eurasians and Neanderthals (D(Y, E; N)) matched estimates from real data (16) (Supplementary note S1a). Our simulation results showed that the estimated dates of Neanderthal gene flow were unbiased when the admixture occurred between 100-1,500 generations ago. However, for dates ≥2000 generations ago, our method underestimated the true dates. Complex demographic events could also lead to a bias in case of older admixtures. Notably, in cases when the population has a history of bottlenecks or expansion events, the inference based on the single-sample statistic was found to be downward-biased for older dates of admixture (≥ 2,500 generations ago) (21). In contrast, even under these complex scenarios, no bias was observed for ancient samples for which the dates of Neanderthal introgression occurred more recently than 1,500 generations ago (21).

To overcome the bias observed at older dates of Neanderthal admixture as expected in extant samples, we applied the admixture LD statistic from (16) (referred to as population-sample statistic). Because the population-sample statistic computes covariance across each pair of markers, it requires data from more than one diploid genome and hence is not applicable to single ancient genomes. Hence, we restricted our analysis with the single-sample statistic to ancient genomes where dates of Neanderthal gene flow were expected to be less than 1,500 generations. We verified via simulations that the application of the population-sample statistic removes the bias observed in (21) and that together these statistics provide accurate estimates (Supplementary note S1b).

To access the utility of our method for estimating the age of ancient genomes (and not just dating Neanderthal gene flow), we simulated data for Neanderthals, west Africans, extant Eurasians and ancient Eurasians (where the individual was sampled between 500-1,750 generations before present). We set the date of shared Neanderthal gene flow to be 2,000 generations before present and Neandertal ancestry proportion in Eurasians to 3% (16, 21). Because of branch shortening, the date of Neanderthal admixture is more recent in ancient than in extant samples. Our simulations showed that the estimated ages of ancient genomes were accurate and that, as expected, the dates were more precise for older samples (Supplementary note S1c).

### Accounting for uncertainty in parameters in real data

Our simulations relied on the accurate modeling of the recombination rate across the genome for dating Neanderthal ancestry. In applications to real data, we used the “shared” African American genetic map referred to as ‘S map’ in (14). This map is inferred by combining information from the deCODE pedigree map in Europeans (based on 500,000 crossovers identified in 15,000 Icelandic meiosis (23)) and the African American genetic map (based on 2.1 million crossovers detected using ancestry switch points observed between African and European ancestry in 30,000 unrelated African Americans) (14). The *S map,* which focuses on the part of the landscape of recombination in African Americans that is shared with Europeans, is known to be the most accurate genetic map available for Eurasians (14).

Despite the high resolution, available maps are less reliable at the short distances (i.e., at kilobase scales) that might be relevant for our analysis. Notably, Sankararaman et al. 2012 showed that the errors in the genetic map can bias the dates of Neanderthal gene flow (16). In order to minimize the bias in estimating ages, we applied the Bayesian “genetic map correction” developed by (16). This correction is a function of the date of Neanderthal gene flow (λ) estimated using a given genetic map and a scalar parameter (*α*) that is related to the precision of the genetic map (larger values of *α* indicate a more precise map). Specifically, *α* is estimated by comparing the genetic distances between a pair of markers as estimated by the *S map* to the crossover distribution observed in a pedigree dataset, such as the deCODE pedigree dataset used here, which is based on 71,929 meiosis (24).

Using this approach, we estimated that the *α* for the *S map* has a value of 3,109 ± 308 per Morgan (M) (see Methods). The effect of this level of map uncertainty is likely to be minimal for ancient samples, in which the ancestry covariance extends to larger distances (greater than hundreds of kilobases). In contrast, for extant samples where the blocks are an order of magnitude smaller, the resulting bias can be substantial, as shown in (16). Thus, we applied the map correction separately for ancient and extant samples to obtain corrected dates of Neanderthal gene flow in generations (*t*_*n*_ in generations).

Further, to convert the dates of gene flow from generations to years (*t*_*n*_ in years) while accounting for some uncertainty in the generation time, we assumed a uniform prior probability distribution on the generation time between 27 and 30 years (25, 26). Mean generation time in ancient humans is not known, but (25) showed that generation times among diverse extant hunter-gatherer and industrialized societies are similar. The difference in the dates of Neanderthal gene flow in ancient and extant genomes then provides an estimate of the age of the ancient genome (*t*_*c*_).

### Case studies

To illustrate the utility of our method, we applied our inference framework to all available ancient modern humans for which whole genome data are available and for which the radiocarbon date is >10,000 years. This threshold was chosen so that the estimated dates of Neanderthal admixture in the ancient genomes are less than 1,500 generations (thus not affected by the bias observed in simulations) and that the difference between the dates of extant and ancient samples is significant (beyond statistic error). We broadly matched the ancestry of the ancient and extant samples, comparing ancient Eurasian samples to 1000 genomes European samples (CEU) (27). We first ascertained markers that were informative for Neanderthal ancestry, i.e., considered sites where the Altai Neanderthal (22) carries the derived allele (relative to the 1000 genomes reconstruction of the human ancestral allele (27)) and 1000 genomes sub-Saharan African samples from Yoruba (YRI) and Luhya (LWK) are homozygous for the ancestral allele (henceforth referred to as ‘ascertainment 0’).

Using the population-sample statistic and genetic map correction described above, we estimated that the Neanderthal admixture in CEU occurred between 1,569– 1,700 generations or 43,189–49,985 yr BP (95% credible interval). This is within the previously published estimate of 37,000–86,000 yr BP (most likely range of 47,000– 65,000 yr BP) based on a different ascertainment scheme, generation time interval and genetic map correction (16). The broader confidence interval in (16) is extremely conservative, reflecting an attempt to take into account biases observed in simulations of complex demographic scenarios. Our inferred date range is narrower as we do not include an additional correction for the effect of model misspecification and we allow for less uncertainty in generation interval compared to (16). Our simulations indicate that the ascertainment 0 should not lead to a bias in dates of Neanderthal gene flow under the complex demographic scenarios applicable to Eurasians, and hence we believe that the additional bias correction is over-conservative (Supplementary Note S1). If our assumptions are valid, dates in the range of 43,189–49,985 years ago are important because they suggest that the main Neanderthal interbreeding with modern humans may have occurred in the context of the Upper Paleolithic expansion of modern humans, rather than at earlier times (28).

We applied our method to estimate the age of five ancient samples: Clovis (29), Mal’ta (30), Kostenki (31), Ust’-Ishim (21) and Oase (32). Below we discuss the dating results for each sample and in Table 1 present a summary of all dates (including estimated ages in generations).

**Table 1:** Estimated ages of ancient genomes

| Sample | Calibrated radiocarbon dates (calBP) | genetic age in generations (mean ± SE) | genetic age in years (mean ± SE) |
| --- | --- | --- | --- |
| Clovis | 12,509 – 12,722 | 624 ± 54 | 17,779 ± 2402 |
| Mal'ta | 23,891 – 24,423 | 862 ± 49 | 24,548 ± 2272 |
| Kostenki | 36,262 – 38,684 | 1,424 ± 43 | 40,544 ± 2026 |
| Ust'-Ishim<sup>a</sup> | 43,210 – 46,880 | 1,373 ± 46 | 39,090 ± 2078 |
| Ust'-Ishim<sup>b</sup> | 43,210 – 46,880 | 1,541 ± 37 | 43,872 ± 1799 |
| Ust'-Ishim<sup>c</sup> | 43,210 – 46,880 | 1,403 ± 38 | 39,974 ± 1823 |
| Ust'-Ishim<sup>d</sup> | 43,210 – 46,880 | 1,590 ± 37 | 45,258 ± 1797 |
| Oase<sup>e</sup> | 37,000 – 42,000 | 1,341 ± 28 | 41,808 ± 1610 |
| Oase<sup>f</sup> | 37,000 – 42,000 | 1,279 ± 30 | 40,052 ± 1647 |
| Oase<sup>g</sup> | 37,000 – 42,000 | 1,343 ± 28 | 41,870 ± 1610 |

Note: Markers were selected based on ascertainment 0, the genetic distances were based on the *S map*, and the genetic correction was based on deCODE data (see Methods).
<sup>a-d</sup> Ust'-Ishim – we tried different models for Ust'-Ishim dating (see results and Figure 2). (A) Dates based on single exponential fit up to genetic distance of 1cM, (B) Dates based on single exponential fit up to genetic distance of 10cM, (C, D) Dates based on double exponential fit ( $y = Ae^{-d\lambda_1} + Be^{-d\lambda_2} + c$ ) to up to genetic distance of 10cM.
<sup>e-g</sup> Oase – we tried different models for Oase dating (see results and Figure 2). (E) Dates based on single exponential fit up to genetic distance of 65cM, (F, G) Dates based on double exponential fit to up to genetic distance of 65cM.

#### Clovis

The Clovis genome from North America has been sequenced to an average coverage of 14.0x (29). The estimated radiocarbon date for this individual is 10,705 ± 35 radiocarbon years (^14^C yr BP) or approximately 12,556–12,707 (95% confidence) calibrated years BP (calBP) (29). As this sample has been sequenced to medium depth coverage, we could not reliably call heterozygous sites and thus restricted analysis to pseudo-diploid genotypes (i.e., we sampled the single majority allele observed in the reads mapped to each site) (see Methods). We applied the single-sample statistic to all pairs of ascertained markers and after applying the genetic map correction estimated that the Neanderthal gene flow in Clovis occurred 28,718 ± 1359 (one standard error) years before he lived. Considering the difference between the dates of Neanderthal admixture in Clovis and CEU provided an estimate of 17,779 ± 2402 years for the age of the Clovis genome. After accounting for uncertainty, this is consistent with its radiocarbon date (Figure 2).

**Figure 2:**
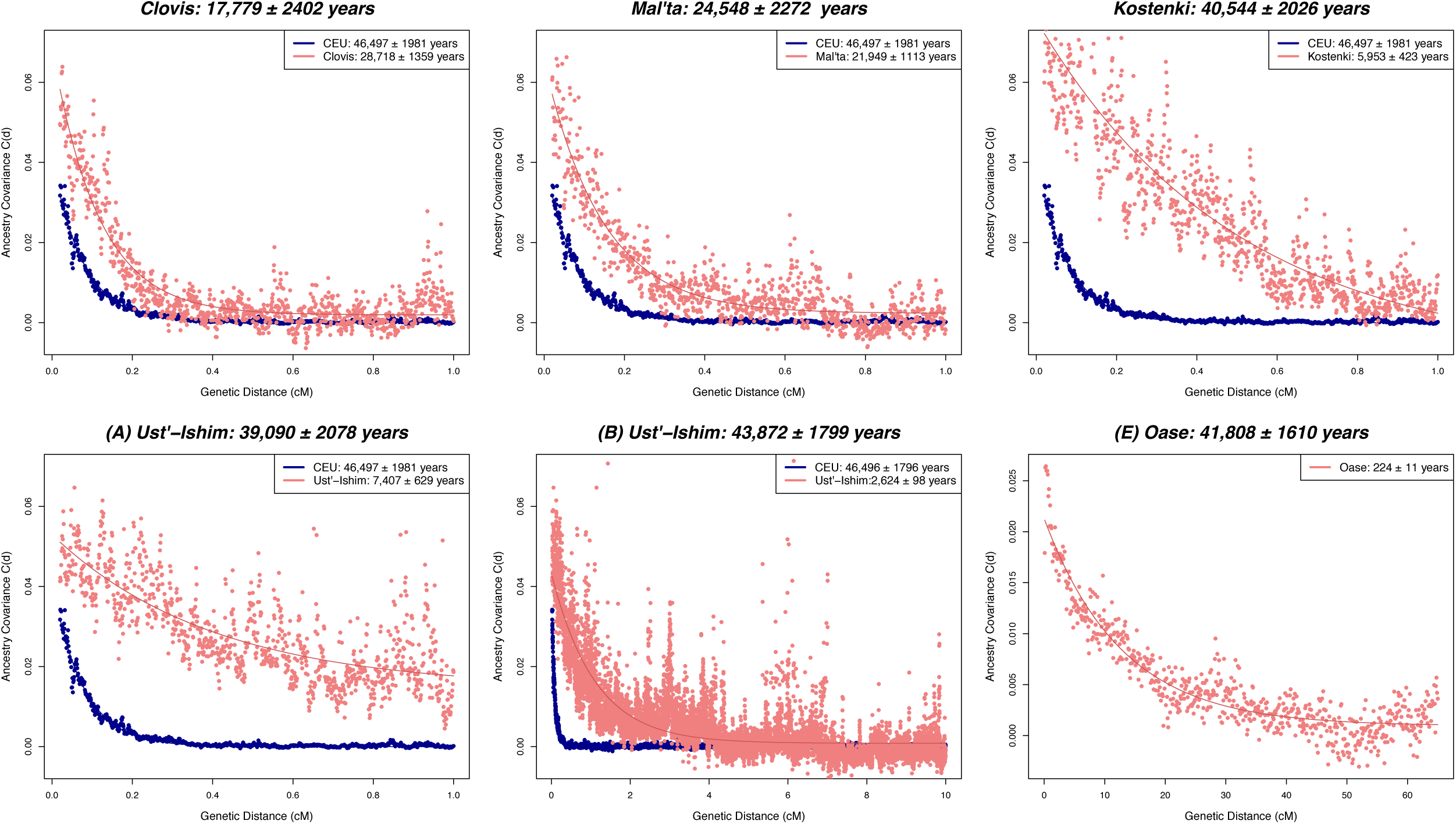
Estimated age of ancient genomes. Estimated dates of Neanderthal gene flow in extant Eurasians (1000 Genomes Europeans (CEU)) shown in blue and ancient Eurasians (either Clovis, Mal’ta, Kostenki, Ust’-Ishim or Oase) shown in pink (see Methods). Estimated ages of the ancient genome (mean ± SE) shown in the title. For Ust’-Ishim, we show two plots: (A) results based on single exponential fit up to the genetic distance of 1cM, (B) results based on single exponential fit up to the genetic distance of 10cM. For Oase, we show results based on single exponential fit up to the genetic distance of 65cM and bin size of 0.1cM. For Oase, we do not show CEU as the analysis was based on a different bin size and maximum distance.

#### Mal’ta

The Mal’ta individual was sampled in south-central Siberia and was sequenced to an average coverage of 1.0x (30). The estimated radiocarbon date of Mal’ta is 20,240 ± 60 ^14^C yr BP or 23,891-24,423 calBP (30). We applied the single-sample statistic to all pairs of ascertained markers, again using pseudo-diploid calls due to the low coverage, and estimated that the Neanderthal gene flow occurred 21,949 ± 1113 years before he lived. In turn, this translates into an estimated age of 24,548 ± 2272 years, similar to the radiocarbon date of this sample (Figure 2).

#### Kostenki

The Kostenki (K14) genome from European Russia has been sequenced to an average coverage of 2.8x (31). The radiocarbon date of this genome is 33,250 ± 500 14C yr BP or 36,262–38,684 calBP (95% confidence) (31). Applying our inference procedure, we estimated that the Neanderthal gene flow in K14 occurred 5,953 ± 423 years before he lived. This is not consistent with recently published estimate of ∼15,000 years before he lived (31). However, details of the genetic map and map correction used in (31) are unclear and could have had a substantial effect on the absolute dates. Considering the difference between the dates of Neanderthal gene flow into K14 and CEU provided an estimated age of 40,544 ± 2026 years (Figure 2), which is consistent with its radiocarbon date.

#### Ust’-Ishim

The Ust’-Ishim genome from western Siberia was sequenced to 42-fold coverage and has been dated twice by ^14^C to be ∼43,210–46,880 calBP (95% confidence) (21). We applied the single-sample statistic to all pairs of ascertained markers, considering diploid genotypes, as the coverage for this sample is high enough to make reliable heterozygous calls. The dates of Neanderthal gene flow in Ust’-Ishim computed based on diploid and pseudo-diploid calls are very similar (Table S6 and discussed later). Applying the single-sample statistic, we estimated the date of the Neanderthal admixture to be 7,407 ± 629 years before he lived, consistent with the date reported in (21) (which used a different genetic map and did not correct for genetic map errors). Considering the difference between the dates of Neanderthal admixture in Ust’- Ishim and CEU, provided an age estimate of 39,090 ± 2078 years, which is consistent with its radiocarbon date (Figure 2).

We note that unlike other ancient genomes, this sample contains many Neanderthal segments that are longer than 1cM, which are poorly fit by the exponential distribution (as the intercept at 1cM is substantially greater than 0) (Figure 2). A likely explanation for the presence of these unmodeled segments is that Ust’-Ishim may not have received all its Neanderthal ancestry from a single pulse of gene flow—or even the same event that affected the extant Eurasian populations, as assumed by our branch shortening model (Figure 1). To explore this possibility, we reran our analysis up to longer distances. Applying our analysis up to 10cM (where the intercept of exponential is almost 0) for Ust’-Ishim and CEU, we estimated that the date of Neanderthal gene flow was 2,624 ± 98 years before Ust’-Ishim lived and 46,496 ± 1796 yr BP respectively. The difference translates to an estimated age of 43,872 ± 1799 for the Ust’-Ishim genome (Figure 2), which is also consistent with the radiocarbon date of the specimen.

The inference of different dates of Neanderthal gene flow in Ust’-Ishim depending on the genetic distance threshold used (unlike in CEU) is consistent with our hypothesis that there may have been at least two pulses of Neanderthal ancestry into the ancestors of Ust’-Ishim, with the curve fitted up to 1cM mostly sensitive to the older events and the curve fitted up 10cM more sensitive to the recent events. To formally test this hypothesis, we applied a likelihood ratio test described in (33) to distinguish between a model of single Neanderthal gene flow and two Neanderthal gene flow events. This provided overwhelming support for the two-pulse model of Neanderthal admixture (p < 10^−20^). Indeed, visualization of the putative Neanderthal ancestry blocks across the Ust’-Ishim genome produces a pattern that is broadly bimodal, with some regions containing >5-10 Mb long blocks (21), which would not be expected unless some of the Neanderthal gene flow occurred recently.

By explicitly fitting a model of two Neanderthal gene flow events, we estimated that the admixture occurred 6,522 ± 312 years and 1,238 ± 58 years before this individual lived (Figure S3). Since it was not clear which of these events might be shared with extant Eurasians, we estimated the age of Ust’-Ishim genome based on each of the two admixture events separately, obtaining 39,974 ± 1823 and 45,258 ± 1797 years (Figure S3). Both of these age estimates are consistent with the radiocarbon date of this sample.

#### Oase

The age of the Oase (Oase 1) genome from Romania has been estimated to be ∼37,000-42,000 calBP by direct radiocarbon dating (32). As the specimen contained tiny amounts of human DNA, it was not feasible to whole genome sequence this individual. Instead, this sample was captured on panels of known markers. The *Archaic panel* (Panel 4, Neanderthal subset: 954,849 markers), where at least one Neanderthal allele differs from the majority allele in a panel of 24 West African Yoruba samples (32), is the most relevant for our application of Neanderthal dating. For this ascertainment, we obtained genotypes for 375,191 markers for CEU and 78,055 for Oase. Using the population-sample statistic, we estimated that the Neanderthal gene flow in CEU occurred 38,410 ± 1477 yr BP, which is lower than the dates based on ascertainment 0. This might be because the *Archaic panel* includes markers where Yoruba is derived and archaic samples contain the ancestral allele. Such sites will likely amplify some background LD, biasing the dates of Neanderthal admixture. Indeed, removal of markers that are polymorphic in Yoruba and ancestral in Altai provides a Neanderthal date estimate of 42,032 ± 1610 yr BP, which is consistent with ascertainment 0. Based on the recommendation in (32), we ran our single-sample statistic up to 65cM for Oase (where the intercept of exponential is almost 0) (see Methods). We estimated that the Neanderthal gene flow in Oase occurred 224 ± 11 years before he lived, consistent with the estimates in (32). Considering the difference with CEU provided an estimated age of 41,808 ± 1610 years, which is consistent with the radiocarbon date of this specimen.

The Oase individual, like Ust’-Ishim, has a bimodal distribution of segments of Neanderthal ancestry. Applying the likelihood ratio test to distinguish between models of single vs. double Neanderthal gene flow events provided strong support for the two-pulse model of Neanderthal admixture (p < 10^−12^). By explicitly fitting a model of two Neanderthal gene flow events, we estimated that the admixture occurred 1,980 ± 348 years and 162 ± 6 years before he lived. This translates into an age estimate of 40,052 ± 1647 and 41,870 ± 1610 years respectively (Figure S3); both of these are consistent with the radiocarbon dates of this specimen.

To provide further confidence that our restriction to markers on the *Archaic panel* provides reliable estimates, we re-estimated the age of Kostenki and Ust’-Ishim that were captured for the same set of markers as Oase. We estimated the age of Kostenki and Ust’-Ishim (maximum distance of 10cM as described earlier) as 39,236 ± 1916 and 38,517 ± 1623 years respectively, consistent with our whole-genome based estimates.

### Robustness

A number of confounding factors could potentially bias genetic age estimates, including uncertainty of genotype calls in low coverage data, our ascertainment of markers, assumptions about the genetic map and effects of natural selection. Below we test how our inferences are affected by each of these factors. For these analyses, we do not include Oase as this individual was only captured on a reduced panel of markers.

#### Robustness to how genotypes were determined

For most of our ancient samples, we did not have sufficient coverage to make reliable diploid calls. Instead, we used pseudo-diploid calls based on the majority allele observed in reads for each sample at each site (see Methods). To verify the robustness of our inferences, we repeated our analysis with pseudo-diploid calls where we sampled a random allele seen in the reads mapped to each site in each sample (pseudo-diploid (random)). In addition, for our high coverage Ust’-Ishim genome, we compared inferences based on diploid and pseudo-diploid calls (both majority and random sampling). For all ancient genomes, the dates were consistent across different genotype calling approaches (Table S6, Figure S2). These results also highlight a strength of our method: that it works well even using pseudo-diploid calls and for samples with low coverage (such as Mal’ta with an average coverage of 1.0x).

#### Robustness to the way markers were selected

To study the effect of marker ascertainment, we analyzed data for additional marker selection schemes shown to be informative for dating Neanderthal ancestry in (16). As previous studies have shown that Luhya have some recent West Eurasian ancestry (∼2.4%) (34), which could affect the dating, we explored an ascertainment of selecting markers only using YRI and Altai Neanderthals (similar to ascertainment used for Oase). The alternative ascertainment schemes considered were ones in which:

(Ascertainment 0) This is presented in the previous section, and is the main ascertainment we use in the study (Figure 2, Table 1). This ascertainment focuses on sites where Neanderthals carry at least one derived allele and all individuals in a panel of sub-Saharan Africans (1000 genomes YRI and LWK) carry the ancestral allele.

(Ascertainment 1) Altai Neanderthal carries the derived allele, Africans (1000 genomes YRI and LWK) carry the ancestral allele, and Eurasians (1000 genomes CEU) are polymorphic.

(Ascertainment 2) Altai Neanderthal carries the derived allele and Eurasians (1000 genomes CEU) are polymorphic and have derived allele frequency of less than 20%. Our threshold is higher than the 10% used in (16). The 20% threshold provides more markers (thus improved precision) and is still informative for Neanderthal ancestry (16).

(Ascertainment 3) Altai Neanderthal carries the derived allele and Africans (1000 genomes YRI) carry the ancestral allele.

The age estimates based on different ascertainments remain consistent for all four ancient genomes and are similar to the radiocarbon dates (Table S7). However, ascertainments 1 and 2 are not symmetric with regard to ancient and extant genomes (extant Eurasians were used for selecting markers while ancient samples were excluded due to their low coverage). This asymmetry could in theory lead to a bias in case the two populations do not share the same admixture history. Thus, ascertainment 3 and ascertainment 0 seem preferable on first principles.

#### Robustness to the genetic map used

A central assumption of our method is that recombination has not changed over time and across populations. Recombination rates, however, are known to have evolved over the course of human evolution evidenced by the observation that the alleles of *PRDM9*, that are the major determinant of recombination hotspots in humans, are changing rapidly (35, 36). Present-day human hotspots appear to have been active for ∼10% of time since divergence from chimpanzees (∼650,000-1.3 million years) (37), suggesting that our assumption is likely valid over the time scale of interest here. Nonetheless, some variation in hotspot usage is known to exist across human populations that split ∼50,000-100,000 years ago (14, 35, 38). Ideally, then, our analysis should be based on a map generated separately for each population. Because such data are unavailable and unlikely to become available for ancient samples, we verified that our inferences are robust to the choice of existing map by repeating the analysis with an African-American map (14) that includes historical hotspots in Africans as well as shared hotspots between Africans and Europeans, and the Oxford CEU LD map (39) that reflects historical recombination rates over tens of thousands of years. We estimated that the map precision (*α*) for these maps as 2,802 ± 14 and 2,620 ± 9 respectively (see Methods), suggesting that these maps are less reliable than the *S map*. This is also reflected in the fact that there is a fair amount of variation in dates across maps. Nonetheless, the age estimates based on the three different maps are consistent (Table S8).

#### Biases due to natural selection

Previous studies have shown that Neanderthal ancestry varies across chromosomes, with unexpectedly large regions devoid of any Neanderthal ancestry, implying a role for natural selection in removing Neanderthal derived alleles from the modern human gene pool (40, 41). In addition, it has been observed that Neanderthal ancestry proportion across the genome is correlated to the B-statistic, a measure of the strength of background selection (40, 42). The B-statistic or B-score measures the reduction in diversity levels at a site due to purifying selection at linked sites, relative to what is expected under strict neutrality (42, 43). We investigated the possibility that natural selection biases our age estimates in two ways: (a) by removing all ascertained markers in regions that are direct targets of natural selection, including conserved elements across primates or coding regions in humans (44), and (b) by studying the variation in estimated ages as a function of the B-score by performing a Block Jackknife analysis (45). We divide the genome in deciles of B-scores as estimated based on the statistic described in (42). Next, we estimated the dates of Neanderthal admixture removing all ascertained markers with a particular B-score and studied the variation in age estimates. In both cases, we found that within the limits of our resolution, the effects of selection on the age estimates was negligible, as the age estimates were consistent after removing putative targets of selection (Table S9) and the Spearman’s correlation between B-statistic and age estimates for all four ancient genomes was not significant (p < 0.05) (Figure S4).

## Discussion

We have developed a novel genetic approach for dating ancient human samples that is applicable for dating ancient non-African genomes that share a history of Neanderthal admixture with extant non-Africans, in the time range of 10,000-45,000 years before present. This method uses the recombination clock as a tool for dating. By characterizing the dates of Neanderthal gene flow in ancient and extant genomes and quantifying the difference in the dates, we infer the age of the ancient genome. Using simulations (Table S5) and real data (Table 1), we show that our method provides reliable age estimates and is applicable for dating low coverage samples such as Mal’ta (average coverage = 1.0x) and Kostenki (average coverage = 2.8x). Further, we find that our inference is robust to marker selection, genetic map errors and natural selection.

Our genetic age estimates take into account a number of sources of uncertainty, including the dates of Neanderthal gene flow, genetic map and generation time. Accounting for the uncertainty in generation time and genetic map has a substantial effect on our standard errors (Table 1). As more data (from pedigrees, genealogies, etc.) and more sensitive methods for inferring historical generation times and recombination rates become available, the estimates of genetic ages will become more precise.

### Comparison with radiocarbon dating

We show that our results are consistent with radiocarbon dates for all studied fossils. While radiocarbon dates are in general more precise than genetic age estimates, our method is complementary to ^14^C dating in that it uses the molecular clock and does not rely on the decay rates of atmospheric isotopes. Thus, our approach provides independent information for the construction and identification of errors in the archaeological records over the past 50,000 years.

A major limitation of our method is that it is not applicable for dating samples that do not share a history of Neanderthal gene flow with non-Africans, such as sub-Saharan Africans. Unlike ^14^C dating the genetic method is unstable for very young samples that are less than 10,000 years old. This is because for a single genome with an old admixture date, it is hard to reliably identify very short segments of Neanderthal ancestry. Over this time range, however, radiocarbon dating is most accurate as calibration can be carried out directly using tree-ring data (46). In contrast, the genetic method becomes more accurate for older samples, as evidenced by our simulations and analysis of real data. In our age estimates the main of source of uncertainty derives from the higher errors in the dates of Neanderthal gene flow in extant genomes.

### Outlook

While in this study we have focused on Neanderthal admixture as our calibration point for dating, there is nothing unique about Neanderthal admixture, and in fact, other shared LD-generating events such as other introgression events (for e.g. the Denisova admixture into Southeast Asians and Oceanians (47)) or other founder events (for e.g. Out of Africa migration (1)) could be used as alternative calibration points through extensions of the methodology reported here. Importantly, if one were to use an older calibration point than the date of Neanderthal gene flow, it could be possible to use genetic data to estimate dates for skeletal remains that are beyond the limits of radiocarbon dating but for which sequence data exists, such as the Altai Neanderthal (22), the Mezmaiskaya Neanderthal (22), or Denisova (48) or for samples that are too small or too old to have enough preserved carbon for radiocarbon dating (like Denisova (48)).

In summary, we have introduced a method that uses genetic data to date ancient human genomes. This method is applicable for dating ancient non-African genomes that are more recent than the Neanderthal gene flow and thus is complementary to ^14^C dating, which interrogates a similar time range using a completely different clock. A notable feature of our method is that it estimates dates in generations (as it is based on recombination), whereas ^14^C dates are determined in years. By comparing radiocarbon dates and branch shortening estimates, it should therefore be possible to obtain estimates of historical generation interval, a key parameter in evolutionary studies.

## Methods and Materials

### Datasets

We used the Altai Neanderthal genome (22) and 1000 genomes Phase I (27) genotypes for sub-Saharan Africans Yoruba (YRI) and Luhya (LWK) for the ascertainment of markers that are informative for dating Neanderthal ancestry. We used the 1000 genomes Phase I northern European samples (CEU) to estimate the date of Neanderthal admixture in extant Eurasians. To identify the ancestral state, we used the 1000 genomes reconstruction of the human ancestral allele, limiting the analysis to positions with confident calls only (27). We applied our inference procedure to all publically available ancient genomes which have radiocarbon dates >10,000 years: Clovis (29), Mal’ta (30), Kostenki (31), Ust’-Ishim (21) and Oase (32). For all ancient samples with whole genome sequence data with less than 30x coverage (Clovis, Mal’ta and Kostenki), we could not reliably call heterozygous genotypes and thus we considered pseudo-diploid majority genotypes, i.e., we chose a single allele at each site for each sample by taking the most common allele (i.e. majority) observed in the reads mapped to that site. When the number of reference and non-reference alleles were the same, we chose a random allele at that site. For testing the robustness of the genotype calling approach, we considered pseudo-diploid calls choosing a random allele from the reads mapped to the site (pseudo-diploid random). For Oase sample, where whole genome sequence data was not available, we used pseudo diploid majority genotype calls for the 78,055 markers from the Archaic panel (panel 4, Neanderthal subset; see (32) for details).

### Statistic for dating Neanderthal admixture

To date Neanderthal mixture, we first ascertained markers that were informative for Neanderthal ancestry. Unless otherwise stated, we used ascertainment 0 where the Altai Neanderthal carries at least one derived allele and 1000 genomes Yoruba and Luhya samples are fixed for the ancestral allele. We did not consider other African populations such as Maasai present in the 1000 Genomes dataset, as recent studies have shown this group has recent ancestry from West Eurasians (34). Moreover, to reduce the effect of sequencing errors, we only considered variant sites where at least one derived allele has been observed among individuals included in the 1000 Genomes Project (except for YRI and LWK). All samples included in the analysis had less than 35% missing data for ascertained markers.

To date Neanderthal admixture in the ancient genomes (Clovis, Mal’ta, Kostenki, Ust’- Ishim and Oase), we used the single-sample statistic and for the extant samples (i.e., the 1000 Genomes CEU), we applied the population-sample statistic (16). Unless otherwise stated, genetic distances were based on the shared African American map (referred to as ‘S’ map in (14)).

### (a) Single-sample statistic

For all pairs of ascertained markers, *S* (*d*)= (*i*, *j*),we computed *C* (*d*), which measures the covariance between genotypes at markers *(i, j*) that are at distance *d* Morgans apart. We ignored markers with missing data.

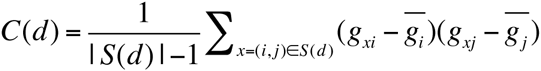

where *g*_*xi*_, *g*_*xi*_ are genotypes at *i*, *j* markers respectively. To estimate the time of mixture (λ), we fitted an exponential distribution with affine (*C*(*d*) = *Ae–d λ*+ *C*) using least squares for genetic distance (*d*) in the range of 0.02 to 1cM in increments of 0.001cM (unless otherwise specified). Because the output based on a single genome is very noisy, the affine term is helpful in capturing some of the noise and is not expected to bias the result.

For Ust’-Ishim, we observed that the intercept at 1cM is substantially greater than 0, hence we also ran the analysis to longer genetic distances, in the range of 0.02 to 10cM in increments of 0.001cM. To estimate the dates of Neanderthal admixture, we tried two models: single exponential (*C*(*d*) = *Ae–d λ*+ *C*) and double exponential (*C*(*d*) = *Ae–d λ*+ *Be–d λ* + *C*)

For Oase, based on the recommendation in (32), we ran our single-sample statistic for genetic distance range of 0.02 to 65cM. As the Neanderthal ancestry blocks are longer in this sample, we used a larger bin size of 0.1cM, which aids in visualization. We note that dates for different bin sizes between 0.001-1cM were consistent (Figure S5). As this sample has a history of multiple Neanderthal gene flow events, we tried two models: single exponential and double exponentials to estimates dates of Neanderthal gene flow.

### (b) Population-sample statistic

For extant populations for which we had access to many individuals, we applied the population-sample statistic described in (16) to estimate the dates of Neanderthal admixture. For all pairs for ascertained markers *(S*(*d*) ) at genetic distance *d*, we computed the statistic:

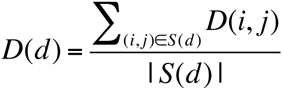

where *D*(*i*, *j*) denotes the standard signed measure of linkage disequilibrium, *D*, at the markers *i*, *j* (17). To estimate the time of mixture (λ), we fitted an exponential distribution (*D*(*d*) =*Ae–d λ*) to this summary statistic using least squares with *d* ranging from 0.02 to 1cM in increments of 0.001cM.

## Genetic map correction and standard error estimation

To account for errors in the genetic map, we applied the genetic map correction described in (16). This approach uses a hierarchical Bayesian model that estimates a scalar parameter *α* that measures the precision of a given genetic map. Specifically, the method compares the expected variance in number of crossovers (based on the number of meiosis in the pedigree dataset) and the observed variance (based on the genetic map) and infers an approximate posterior distribution of *α* by Gibbs sampling. Simulations reported in (16) show that this method is effective in characterizing the map uncertainty for the deCODE and Oxford CEU LD maps (39).

Using this approach, we estimated *α* separately for the *S map*, African American map (16) and Oxford CEU LD map (39) as 3,414 ± 13, 2,802 ± 14 and 2,620 ± 9 per Morgan respectively, by comparing each map to the distribution of cross-overs observed in the deCODE pedigree dataset containing 71,929 meiosis detected in Icelandic pedigrees (24). As discussed in (49), our estimates for the *S map* and Oxford CEU LD map are higher than reported in (16), which used the Oxford CEU LD map and the genetic map correction based on 728 meiosis detected in European American Hutterite pedigrees (50). The finer resolution of the deCODE pedigree dataset used for cross-checking likely explains this discrepancy (49). However, as the *S map* uses a subset of the deCODE pedigree data (15,000 meiosis) as a prior for the construction of the genetic map, and then uses a larger dataset (71,929 meiosis) for estimation of *α*, we believe this could lead to an overestimation of *α* for the *S map*. To address this issue in a conservative way, we combined the information for *α* for the African American map (that does not use the deCODE data as prior) and the *S map*, to estimate the value of *α* for the *S map* as 3,109 ± 308 per Morgan.

Given an estimate of (*α*) and date of Neanderthal gene flow (?) (either estimated using *C(d)* or *D(d)),* we computed the corrected date of gene flow in generations (*t*_*n*_ in generations) using:

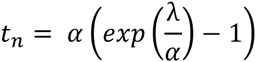

Further, we assumed a uniform prior distribution on the number of years per generation of 27–30 years (51, 52) and integrated over the uncertainty in generation time to obtain corrected dates of gene flow in years (*t*_*n*_ in years).

## Estimating the age of the ancient genomes

To estimate the age (*t*_*c*_) of the ancient genome of interest (Clovis, Mal’ta, Kostenki, Ust’-Ishim and Oase), we quantified the difference in dates of Neanderthal admixture in ancient genome (*t*_*na*_) (estimated using the single-sample statistic) and extant CEU genomes (*t*_*ne*_) (estimated using the population-sample statistic). Standard errors were estimated based on the Bayesian framework described above. We report the mean and standard error of genetic ages in generations and years.

## Web resources

The software implementing the dating approach will be made available for download from URL: http://genetics.med.harvard.edu/reichlab/Reich_Lab/Software.html upon publication of this manuscript.

## Acknowledgements

We thank Mark Lipson for helpful discussions about characterizing the uncertainty in the genetic map. We thank Bridget Alex, Bence Viola, Katerina Douka, Thomas Higham, Svante Pääbo and David Pilbeam for comments on the manuscript. PM was supported by the National Institutes of Health (NIH) under Ruth L. Kirschstein National Research Service Award F32 GM115006-01. DR and NP were supported by U.S. National Science Foundation HOMINID grant BCS-1032255 and U.S. National Institutes of Health grant GM100233. DR is a Howard Hughes Medical Institute Investigator.

## Supplementary Material

### Note S1: Simulations

To assess the performance of our method, we performed coalescent simulations for various demographic scenarios. For all simulations described below, we used the coalescent simulator *ms* (53) to generate data for 50 regions of 50 Mb each for Eurasians, west Africans and Neanderthals with the following parameters. These parameters were chosen to match the F_ST_ (west Africans, Eurasians) and D(west Africans, Eurasians; Neanderthals) observed in real data (Table S1).

- Mutation rate = 1.5x10^−8^ per bp/generation
- Recombination rate = 1x10^−8^ per bp/generation
- Effective population size (Ne) of modern humans = 10,000.
- Effective population size (Ne) of Neanderthals = 2,500.
- Neanderthals were sampled 2,000 generations ago.
- Eurasians split from Africans 3,000 generations before present.
- Modern humans split from Neanderthal 12,000 generation before present.
- Gene flow from Neanderthals into Eurasians occurred instantaneously and contributed 3% ancestry to Eurasians.

*(1) Date of Neanderthal admixture in Eurasians varying in all extant individuals between 100-2,500 generations ago*

We simulated data using *ms* for 44 haploid genomes: 20 Eurasians, 20 west Africans, and four Neanderthals. We simulated data under a simple demographic model with gene flow from Neanderthals into Eurasians that occurred instantaneously at varying times of mixture (100-2,500 generations ago). We sampled Neanderthals 2,000 generations ago; to achieve this, we set the effective population size to be very small (order of 10^−6^) and the sample size of Neanderthals to one chromosome during the time period of 0-2,000 generations. This has the effect of leading to a negligible rate of recombination and mutation during this interval. In addition, to remove all the mutations that occurred during this period, we simulated two chromosomes with similar history and removed any sites that were not shared between these chromosomes. We combined two haploid chromosomes at random to generate 22 diploid chromosomes.

The *ms* command line is as follows:

*ms* 44 1 -r 20000 50000000 -t 30000 -I 6 20 20 1 1 1 1 -en 0 1 1 -en 0 2 1 -en 0 3 1e-10 -en 0 4 1e-10 -en 0 5 1e-10 -en 0 6 1e-10 -es *t*_*n*_ 2 0.97 -en 0.02500025 7 0.25 -en 0.02500025 2 1 -ej 0.05 4 3 -ej 0.05 6 5 -en 0.05000025 3 0.25 -en 0.05000025 5 0.25 -ej 0.0500025 5 3 -en 0.050005 3 0.25 -ej 0.075 2 1 -en 0.0750025 1 1 –ej 0.1 7 3 -en 0.1000025 3 0.25 -ej 0.3 3 1 -en 0.3000025 1 1

Here, pop1 = west Africans, pop2 = Eurasians, pop3 = Neanderthals and *t*_*n*_ = time of Neanderthal gene flow (between 500 – 2,500 generations). Estimated F_ST_ and D-statistics for each simulation are shown in Table S1.

**Table S1:**
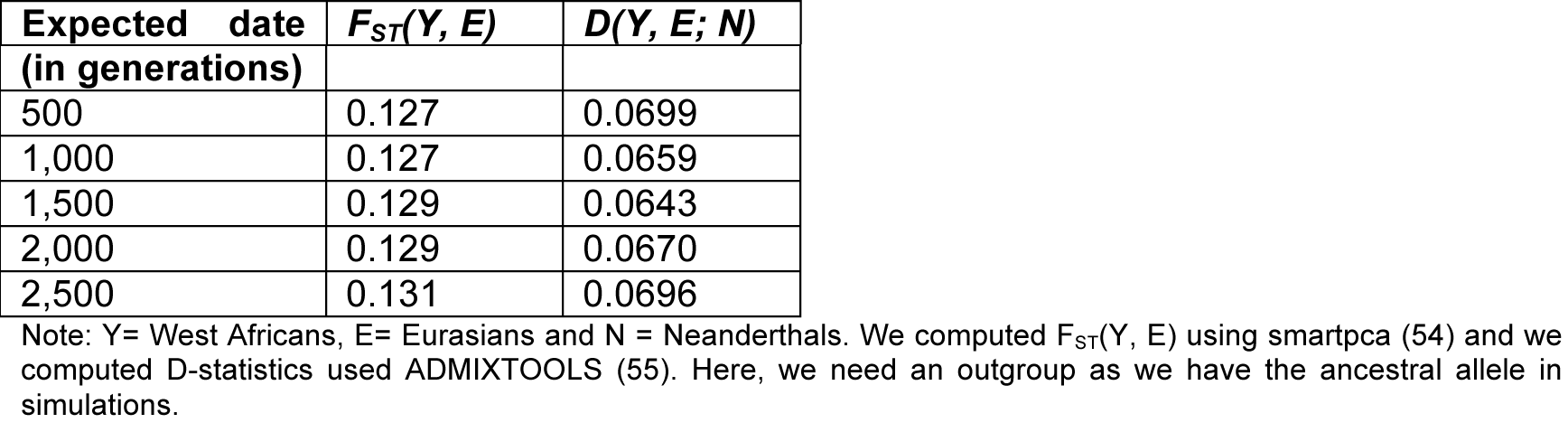
Summary statistics in simulated data.

To date the Neanderthal gene flow in each simulated Eurasian, we ascertained Neanderthal informative markers based on the ascertainment scheme where Neanderthals carry the derived allele and west Africans are fixed for the ancestral allele. We applied the single-sample statistic (see Methods) to all pairs of ascertained markers. We fitted a single exponential distribution to estimate the date of the Neanderthal gene flow. To compute standard errors, we used a weighted block jackknife procedure (45), removing one region (50MB) in each run to estimate the variability in the estimated dates across the genome. Figure S1 shows the results of the simulation. For dates between 100-1,500 generations, we obtained accurate and precise dates of mixture. However, for dates, ≥2000 generations, we observed a downward bias in the estimated dates and decreased precision.

Next, we applied the population-sample statistic (see Methods) to date the Neanderthal admixture in Eurasians. For each date of mixture, we ran the population-sample statistic on five random simulated diploid individuals. For all time-depths, we obtained relatively unbiased dates of mixture (Figure S1).

**Figure S1:**
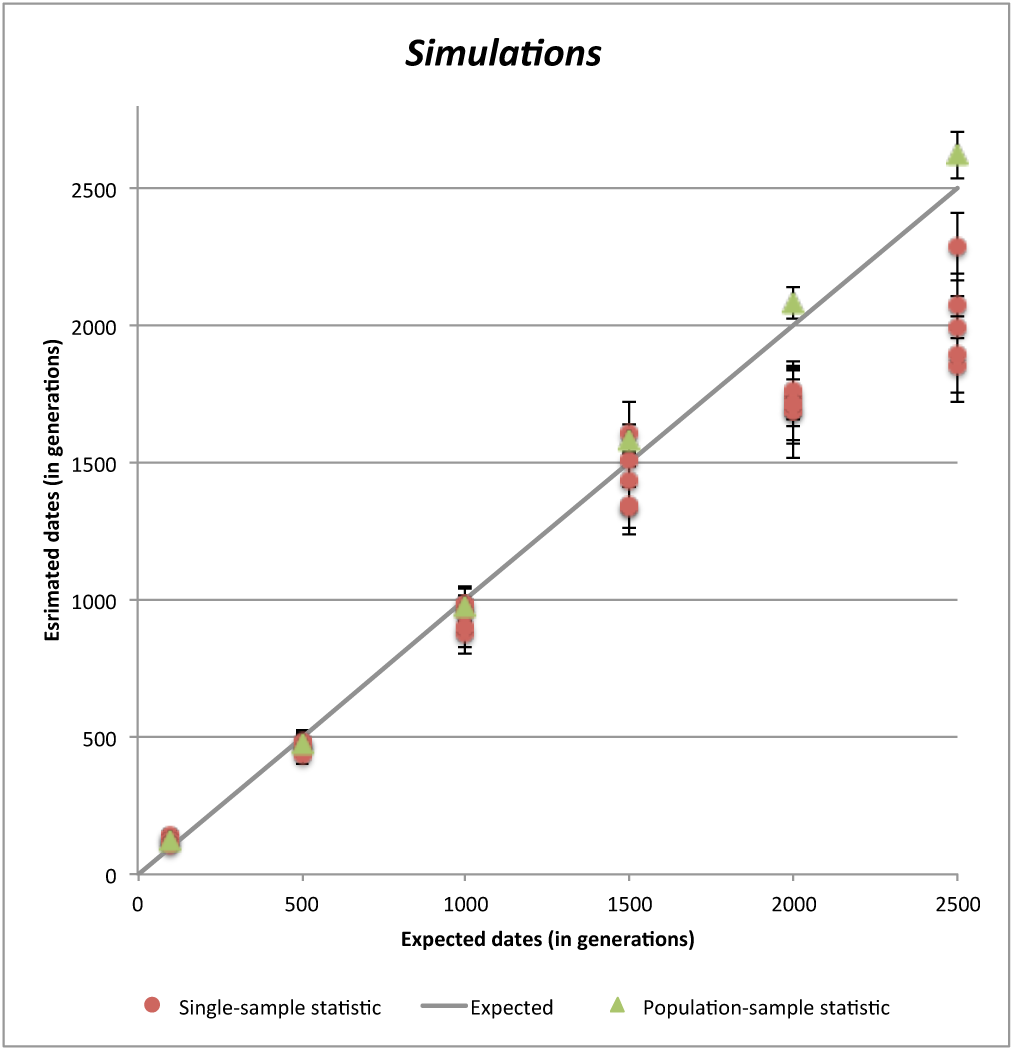
Estimated dates of Neanderthal gene flow. For each time point λ, we simulated ten diploid Eurasian genomes that derive 3% of their ancestry from Neanderthal gene flow that occurred λ generations ago. Shown is the true time against the estimated mean and standard error of the dates based on two statistics (single-sample and population-sample statistics) in generations. Standard errors were computed based on a weighted block jackknife procedure (45), removing one region (50MB) in each run.

*(2) Population-sample statistic removes bias observed in Fu et al. (2014)*

Simulations reported in Fu et al. (2014) showed that under complex scenarios of multiple bottleneck and expansions, the single-sample statistic gives downward biased results for the older dates of Neanderthal admixture (>2000 generations ago). In contrast, little or no bias was observed for ancient samples or recent dates of Neanderthal admixture (21). Here, we used the same data generated by Fu et al. (2014) (Supplementary Information S18: Simulation 4) and applied the population-sample statistic (see Methods). Briefly, Fu et al. (2014) simulated data for 10 African, 20 European (Eu), 20 Asian (As), one Altai Neanderthal and one ancient non-African genome (Ust’-Ishim) based on the demographic model described in Gravel et al. (2011). Detailed parameters for the simulations are described in (21). We note that the observed F_ST_ between west Africans and Eurasians and D-statistics in this simulation are more extreme than observed in real data, likely because some of the parameters (such as bottleneck time and strength) are more extreme than realistic (Table S2).

**Table S2:** Summary statistics in simulated data from Fu et al. (2014).

| Statistic applied | Estimated |
| --- | --- |
| $F_{ST}(Y, E)$ | 0.236 |
| $F_{ST}(Y, A)$ | 0.257 |
| $F_{ST}(E, A)$ | 0.242 |
| $D(Y, E; N)$ | 0.0745 |
| $D(Y, A; N)$ | 0.0663 |

Note: Y= West Africans, E= Europeans, A=Asians and N = Neanderthals.

To estimate the date of Neanderthal gene flow, we applied the single-sample and population-sample statistic to all pairs of ascertained markers. We estimated the dates of Neanderthal admixture by fitting exponential distributions to each of the outputs of the two statistics. Standard errors were computed using a weighted Block Jackknife as described above. Table S3 shows the results of the simulations. While the dates of the Neanderthal admixture based on the single-sample statistic are downward biased, the dates based on the population-sample statistic are accurate.

**Table S3:**
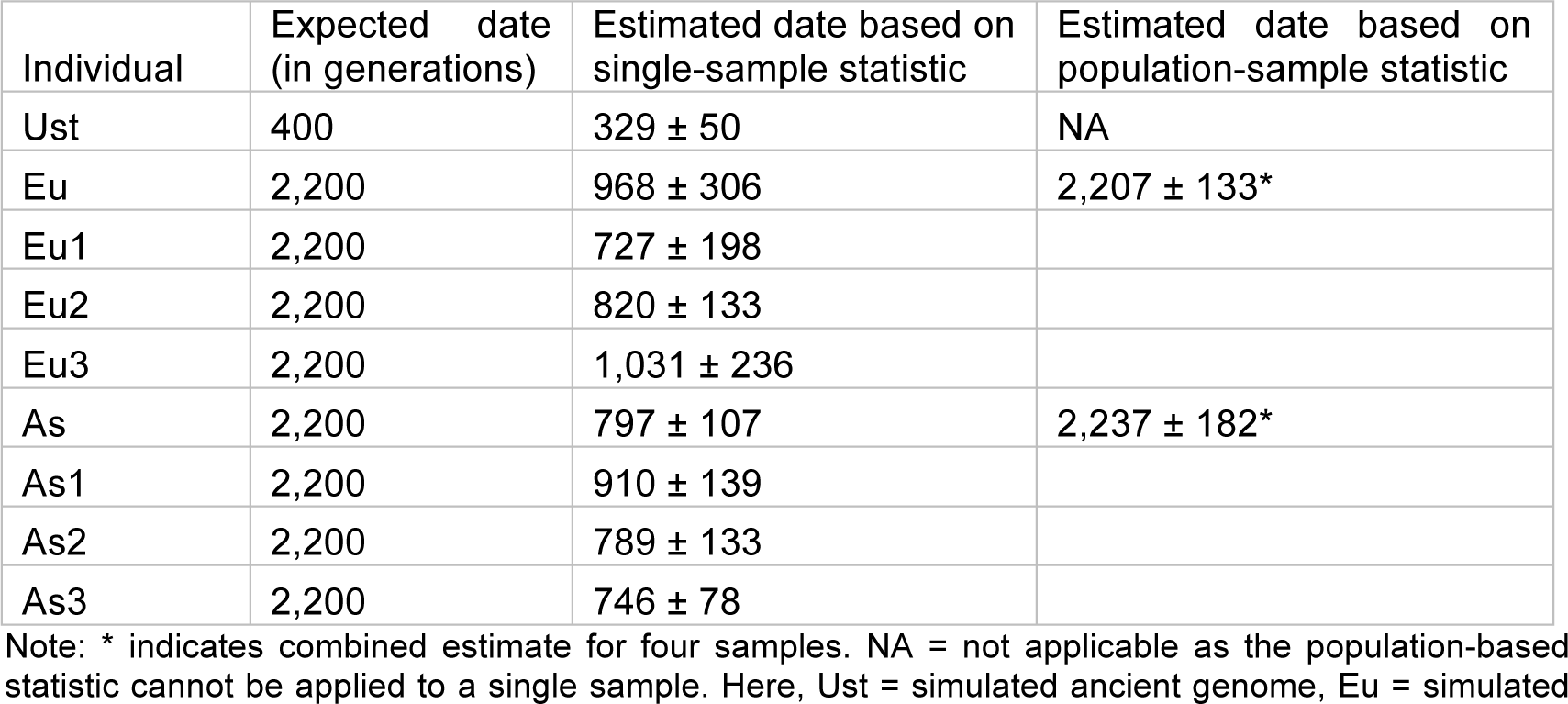
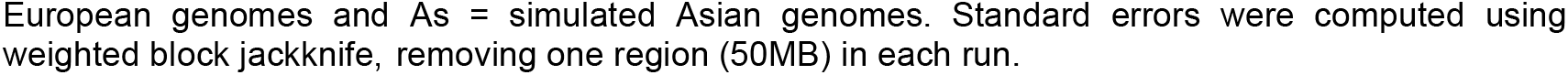
Comparison of single-sample and population-sample statistics for dating Neanderthal admixture.

| Individual | Expected date (in generations) | Estimated date based on single-sample statistic | Estimated date based on population-sample statistic |
| --- | --- | --- | --- |
| Ust | 400 | 329 ± 50 | NA |
| Eu | 2,200 | 968 ± 306 | 2,207 ± 133* |
| Eu1 | 2,200 | 727 ± 198 |  |
| Eu2 | 2,200 | 820 ± 133 |  |
| Eu3 | 2,200 | 1,031 ± 236 |  |
| As | 2,200 | 797 ± 107 | 2,237 ± 182* |
| As1 | 2,200 | 910 ± 139 |  |
| As2 | 2,200 | 789 ± 133 |  |
| As3 | 2,200 | 746 ± 78 |  |

Note: \* indicates combined estimate for four samples. NA = not applicable as the population-based statistic cannot be applied to a single sample. Here, Ust = simulated ancient genome, Eu = simulated
European genomes and As = simulated Asian genomes. Standard errors were computed using weighted block jackknife, removing one region (50MB) in each run.

*(3) Estimating the age of ancient genomes using the Neanderthal ancestry signal*

To verify the reliability of our inference method for estimating the age of an ancient sample, we performed coalescent simulations where we sampled ancient Eurasians at some time in the past (between 500-1,750 generations ago). We simulated these samples in the same way as we simulated Neanderthals, so the sample consists of one chromosome between the present and sampling time. Ancient Eurasians and extant Eurasians share the gene flow from Neanderthals that occurred 2,000 generations ago. The observed F_ST_ and D are as shown in Table S4.

**Table S4:**
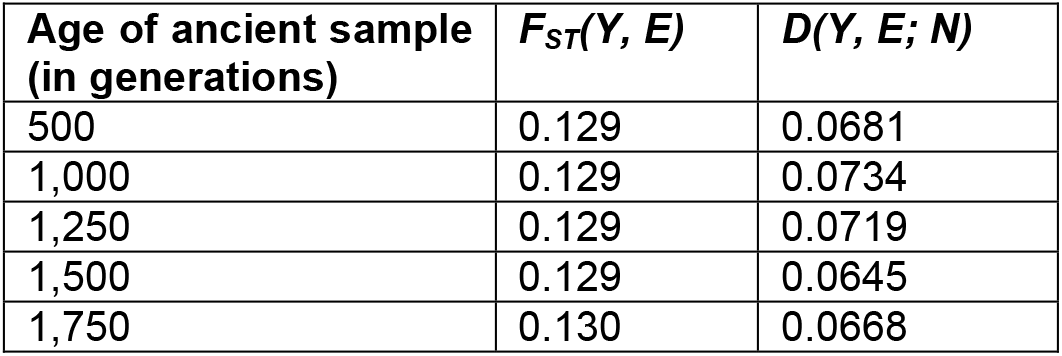
Summary statistics in simulated data.

To estimate the age of the ancient genome, we first estimated the dates of Neanderthal admixture in ancient genomes (using the single-sample statistic) and extant genomes (using the population-sample statistic). The difference in the Neanderthal gene flow dates provides an estimate of the age of the ancient genome. Standard errors were computed using weighted Block Jackknife as described above. The inference procedure based on using the two statistics provides reliable dates for the age of the ancient genomes (Table S5).

**Table S5.** Estimated ages of simulated ancient genomes.

| Sim. | Estimated date of gene<br>flow in extant ($t_{na}$)<br>mean ± SE (Expected) | Estimated date of gene<br>flow in ancient ($t_{ne}$)<br>mean ± SE (Expected) | Estimated age of ancient<br>sample ($t_c$)<br>mean ± SE (Expected) |
| --- | --- | --- | --- |
| Sim 1 | 2060 ± 43<br>(2000) | 1312 ± 123<br>(1500) | 748 ± 130<br>(500) |
| Sim 4 | 2021 ± 37<br>(2000) | 945 ± 71<br>(1000) | 1075 ± 80<br>(1000) |
| Sim 3 | 2116 ± 44<br>(2000) | 762 ± 46<br>(750) | 1354 ± 64<br>(1250) |
| Sim 2 | 2092 ± 44<br>(2000) | 561 ± 38<br>(500) | 1531 ± 58<br>(1500) |
| Sim 5 | 2135 ± 46<br>(2000) | 271 ± 25<br>(250) | 1864 ± 52<br>(1750) |

Note: Expected dates shown in brackets in red. Standard errors were computed using weighted block jackknife, removing one region (50MB) in each run.

**Table S6:** Effect of genotype calling approach.

| Sample | *Diploid* | *Pseudo-diploid (major)* | *Pseudo-diploid (random)* |
| --- | --- | --- | --- |
| Clovis | n/a | 17,779 ± 2402* | 17,168 ± 2364 |
| Mal'ta | n/a | 24,548 ± 2272* | 24,478 ± 2254 |
| Kostenki | n/a | 40,544 ± 2026* | 39,725 ± 2034 |
| Ust'-Ishim (B) | 43,872 ± 1799* | 43,884 ± 1799 | 43,821 ± 1984 |

Note: Markers were selected based on ascertainment 0, the genetic distances were based on the *S map*, and the genetic correction was based on data from (23) (see Methods). n/a – indicates that the coverage is not sufficient to make reliable diploid calls. Ust'-Ishim dates are based on the model of a single exponential fitted to genetic distance of up to 10cM.

**Table S7:** Effect of marker ascertainment on age estimates.

| Sample | *ascertainment 0\** | *ascertainment 1* | *ascertainment 2* | *ascertainment 3* |
| --- | --- | --- | --- | --- |
| Number of ascertained markers included | 221,482 | 164,995 | 486,804 | 280,441 |
| Date of Neanderthal gene flow in CEU | 46,497 ± 1981 | 46,341 ± 1882 | 48,122 ± 2106 | 41,593 ± 1658 |
| Age of Clovis | 17,779 ± 2402 | 19,446 ± 2150 | 17,972 ± 2479 | 12,755 ± 2091 |
| Age of Mal'ta | 24,548 ± 2272 | 23,688 ± 2109 | 19,716 ± 2526 | 22,836 ± 1878 |
| Age of Kostenki | 40,544 ± 2026 | 37,511 ± 1930 | 39,259 ± 2157 | 34,379 ± 1715 |
| Age of Ust'-Ishim (B) | 43,872 ± 1799 | 41,097 ± 1686 | 44,823 ± 1844 | 38,990 ± 1599 |

Note: \* indicates the approach used in main text. For all four ascertainments, we used the genetic distances based on the *S map* and the genetic map correction based on data from (23). Ust'-Ishim dates are based on model of single exponential fitted to genetic distance of up to 10cM.

**Table S8:** Effect of genetic map on age estimates.

| Sample | *S map* * | *AA map* | *Oxford CEU map* |
| --- | --- | --- | --- |
| Date of Neanderthal gene flow in CEU | 46,497 ± 1981 | 41,658 ± 1444 | 42,700 ± 1505 |
| Clovis | 17,779 ± 2402 | 16,452 ± 1876 | 18,798 ± 1915 |
| Mal'ta | 24,548 ± 2272 | 21,935 ± 1675 | 25,940 ± 1755 |
| Kostenki | 40,544 ± 2026 | 35,996 ± 1531 | 37,790 ± 1587 |
| Ust'-Ishim (B) | 43,872 ± 1799 | 38,931 ± 1372 | 39,527 ± 1460 |

Note: \* indicates the approach used in main text. We used ascertainment 0 described in the Methods. Pedigree based maps from deCODE were not used as our genetic map correction is based on these data. Ust'-Ishim dates are based on the model of single exponential fitted to genetic distance of up to 10cM.

**Table S9:** Effect of natural selection on age estimates.

| Sample | *Standard Approach* * | *Remove putative targets of selection* |
| --- | --- | --- |
| Number of ascertained markers included | 221,482 | 213,423 |
| Clovis | 17,779 ± 2402 | 17,451 ± 2381 |
| Mal'ta | 24,548 ± 2272 | 24,331 ± 2244 |
| Kostenki | 40,544 ± 2026 | 40,199 ± 2026 |
| Ust'-Ishim (B) | 43,872 ± 1799 | 43,774 ± 1884 |

Note: \* indicates the approach used in main text. Markers were selected based on ascertainment 0, the genetic distances were based on the *S map*, and the genetic correction was based on data from (23) (see Methods). Ust'-Ishim dates are based on model of single exponential fitted to genetic distance of up to 10cM. To account for putative targets of selection, we remove ascertained markers belonging to exons (http://hgdownload.cse.ucsc.edu/goldenPath/hg19/database/refGene.txt.gz) or conserved elements in primates (http://hgdownload.cse.ucsc.edu/goldenPath/hg19/database/phastConsElements46wayPrimates.txt.gz).

**Figure S2:**
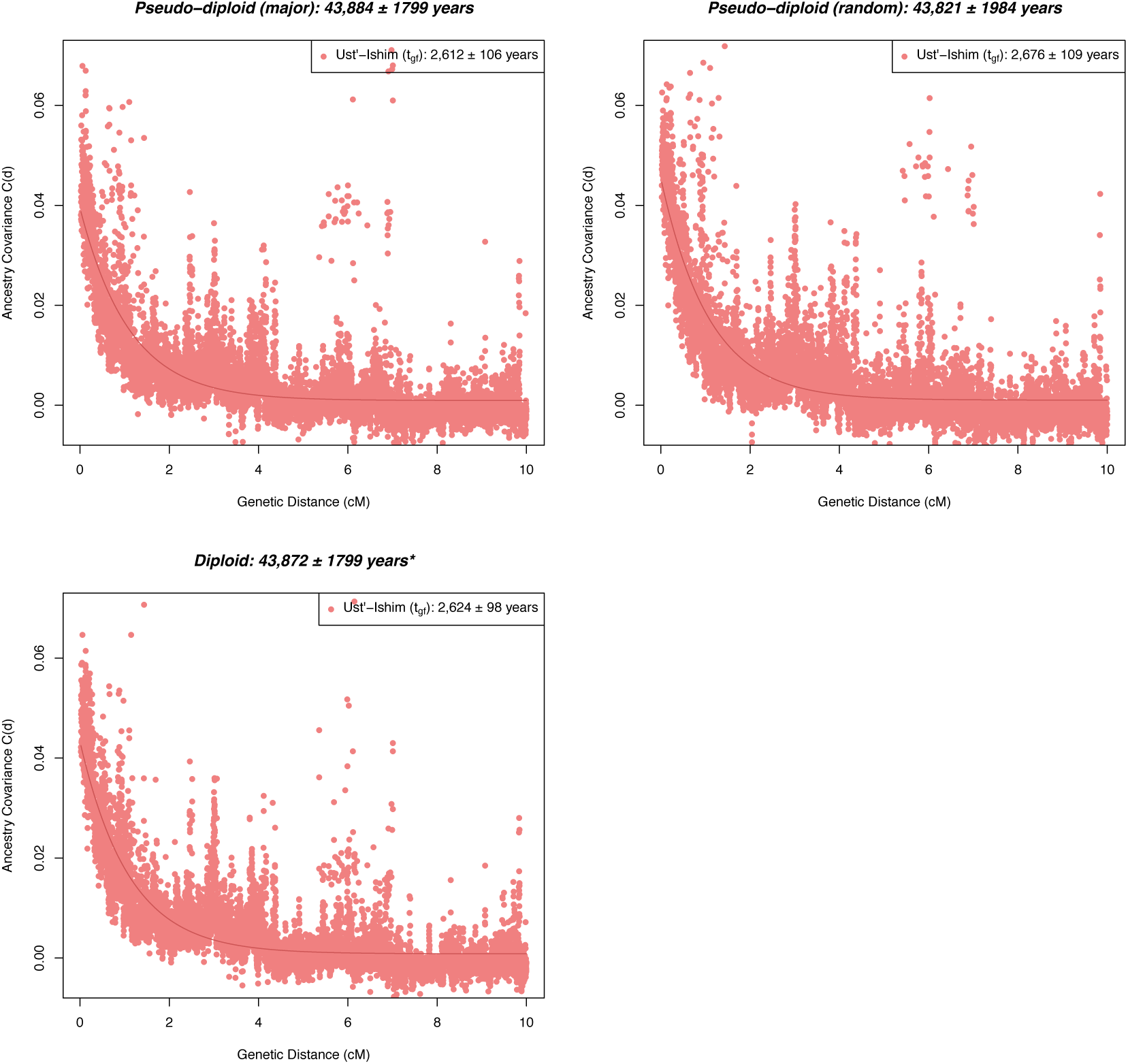
Dates of Neanderthal admixture in Ust’-Ishim based on three approaches of genotype determination. We determined genotypes for markers selected based on ascertainment 0 using (a) Pseudo-diploid majority call, where we chose the major allele in case of heterozygous sites, (b) Pseudo-diploid random call, where we chose a random allele in case of heterozygous sites, and (c) Diploid call, where we considered heterozygous calls. * indicates the results described in the main text.

**Figure S3:**
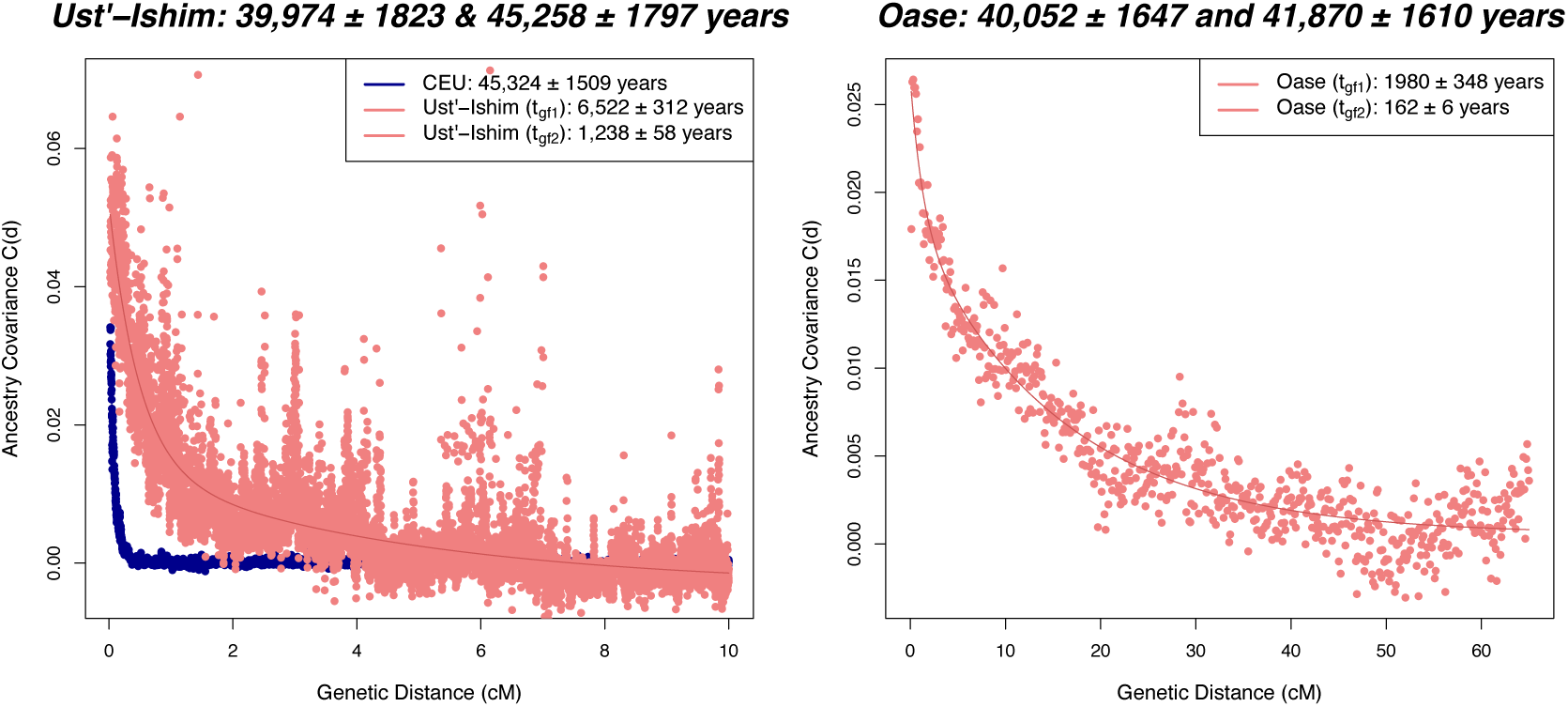
Estimated age of Ust’-Ishim and Oase genomes using a model of two Neandertal admixture events. Estimated dates of Neanderthal gene flow in extant Eurasians (1000 Genomes Europeans (CEU)) shown in blue and ancient Eurasians (either Ust’-Ishim or Oase) shown in pink (see Methods). Estimated ages of the ancient genome (mean ± SE) shown in the title. For Ust’-Ishim, we show results based on double exponential fit up to the genetic distance of 10cM. For Oase, we show results based on double exponential fit up to the genetic distance of 65cM and bin size of 0.1cM. For Oase, we do not show CEU as the analysis was based on a different bin size.

**Figure S4:**
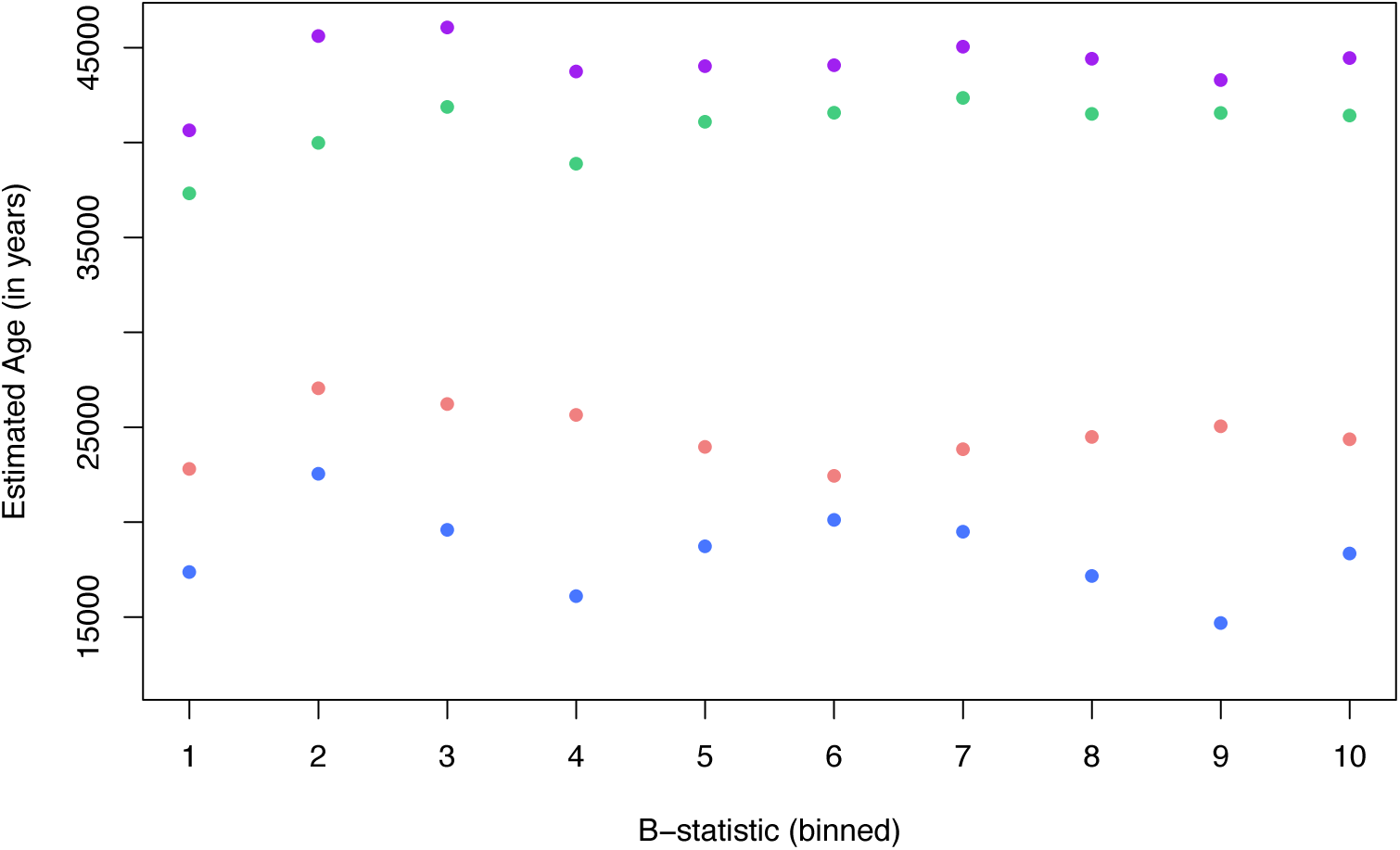
Effect of B-statistic on age estimates. To capture the effect of B-statistics on age estimates, we performed a block jackknife, removing ascertained markers that have a particular value of B-statistic (shown on X axis) in each run and studying the variation in age estimates (Y-axis). Spearman’s rank correlation test showed that the slope is not significant (p < 0.05) for any of the four genomes.

**Figure S5:**
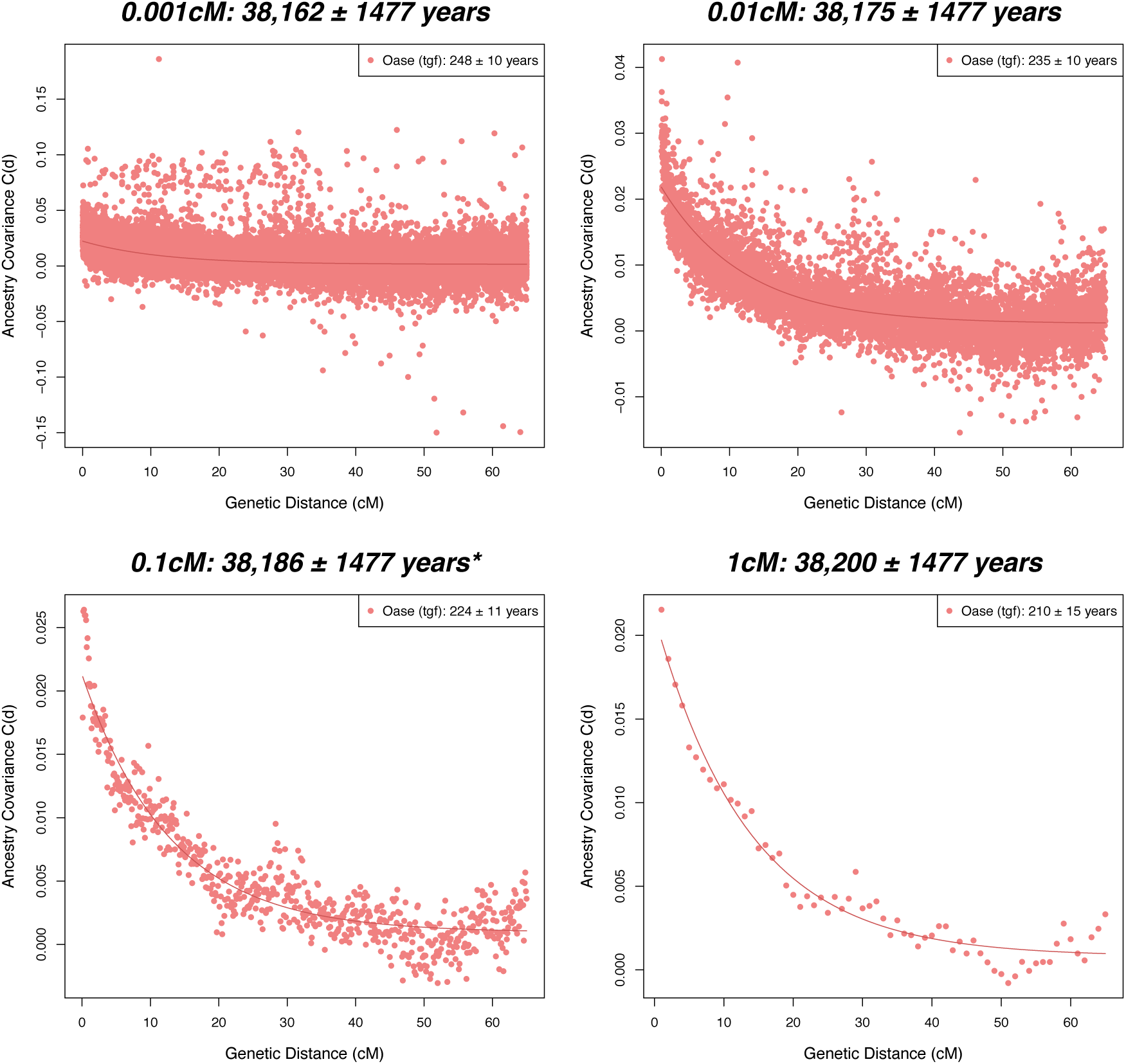
Age estimates for Oase for different bin sizes. As the estimates of Oase are very noisy for a bin size of 0.001cM, we tried larger bin sizes. The title of each sub-figure shows the bin size and the estimated age. In each case, we compared the dates of Neanderthal gene flow with CEU where the Neanderthal gene flow occurred 42,032 ± 1610 yr BP (for this ascertainment). * indicates the estimates used in main text.

## References

1. Pickrell JK & Reich D (2014) Toward a new history and geography of human genes informed by ancient DNA. Trends in Genetics 30(9):377–389.

2. Bronk Ramsey C (2008) Radiocarbon dating: Revolutions in Understanding*. Archaeometry 50(2):249–275.

3. Anderson EC & Libby W (1951) World-wide distribution of natural radiocarbon. Physical Review 81(1):64.

4. Godwin H (1962) Half-life of radiocarbon. Nature 195.

5. Damon PE, Lerman JC, & Long A (1978) Temporal fluctuations of atmospheric 14C: causal factors and implications. Annual Review of Earth and Planetary Sciences 6:457.

6. Aitken MJ (2014) Science-based dating in archaeology (Routledge).

7. Jakobsson M, et al. (2008) Genotype, haplotype and copy-number variation in worldwide human populations. Nature 451(7181):998–1003.

8. Stadler T & Yang Z (2013) Dating phylogenies with sequentially sampled tips. Systematic biology 62(5):674–688.

9. Fu Q, et al. (2013) A revised timescale for human evolution based on ancient mitochondrial genomes. Current Biology 23(7):553–559.

10. Nesheva D (2014) Aspects of Ancient Mitochondrial DNA Analysis in Different Populations for Understanding Human Evolution. science 1(5):6.

11. Meyer M, et al. (2012) A high-coverage genome sequence from an archaic Denisovan individual. Science 338(6104):222–226.

12. Scally A & Durbin R (2012) Revising the human mutation rate: implications for understanding human evolution. Nature Reviews Genetics 13(10):745-753.

13. Ségurel L, Wyman MJ, & Przeworski M (2014) Determinants of mutation rate variation in the human germline. Annual review of genomics and human genetics 15:47–70.

14. Hinch AG, et al. (2011) The landscape of recombination in African Americans. Nature 476(7359):170–175.

15. Green RE, et al. (2010) A draft sequence of the Neandertal genome. science 328(5979):710–722.

16. Sankararaman S, Patterson N, Li H, Pääbo S, & Reich D (2012) The date of interbreeding between Neandertals and modern humans. PLoS Genetics 8(10):e1002947.

17. Chakraborty R & Weiss KM (1988) Admixture as a tool for finding linked genes and detecting that difference from allelic association between loci. Proceedings of the National Academy of Sciences 85(23):9119–9123.

18. Hellenthal G, et al. (2014) A genetic atlas of human admixture history. Science 343(6172):747–751.

19. Moorjani P, et al. (2011) The history of African gene flow into Southern Europeans, Levantines, and Jews. PLoS genetics 7(4):e1001373.

20. Loh P-R, et al. (2013) Inferring Admixture Histories of Human Populations Using Linkage Disequilibrium. Genetics 193(4):1233–1254.

21. Fu Q, et al. (2014) Genome sequence of a 45,000-year-old modern human from western Siberia. Nature 514(7523):445–449.

22. Prüfer K, et al. (2014) The complete genome sequence of a Neanderthal from the Altai Mountains. Nature 505(7481):43–49.

23. Kong A, et al. (2010) Fine-scale recombination rate differences between sexes, populations and individuals. Nature 467(7319):1099–1103.

24. Kong A, et al. (2014) Common and low-frequency variants associated with genome-wide recombination rate. Nature genetics 46(1):11–16.

25. Fenner J (2005) Cross-cultural estimation of the human generation interval for use in genetics-based population divergence studies. American Journal of Physical Anthropology 128(2):415.

26. Helgason A, Hrafnkelsson B, Gulcher JR, Ward R, & Stefv°nsson Kr (2003) A populationwide coalescent analysis of Icelandic matrilineal and patrilineal genealogies: evidence for a faster evolutionary rate of mtDNA lineages than Y chromosomes. The American Journal of Human Genetics 72(6):1370–1388.

27. Consortium GP (2010) A map of human genome variation from population-scale sequencing. Nature 467(7319):1061–1073.

28. Higham T, et al. (2014) The timing and spatiotemporal patterning of Neanderthal disappearance. Nature 512(7514):306–309.

29. Rasmussen M, et al. (2014) The genome of a Late Pleistocene human from a Clovis burial site in western Montana. Nature 506(7487):225–229.

30. Raghavan M, et al. (2013) Upper Palaeolithic Siberian genome reveals dual ancestry of Native Americans. Nature.

31. Seguin-Orlando A, et al. (2014) Genomic structure in Europeans dating back at least 36,200 years. Science 346(6213):1113–1118.

32. Fu Q, et al. (2015) An early modern human from Romania with a recent Neanderthal ancestor. Nature.

33. Moorjani P, et al. (2013) Genetic evidence for recent population mixture in India. American journal of human genetics 93(3):422–438.

34. Pickrell JK, et al. (2014) Ancient west Eurasian ancestry in southern and eastern Africa. Proceedings of the National Academy of Sciences 111(7):2632–2637.

35. Baudat F, et al. (2010) PRDM9 is a major determinant of meiotic recombination hotspots in humans and mice. Science 327(5967):836–840.

36. Myers S, et al. (2010) Drive against hotspot motifs in primates implicates the PRDM9 gene in meiotic recombination. Science 327(5967):876–879.

37. Lesecque Y, Glémin S, Lartillot N, Mouchiroud D, & Duret L (2014) The Red Queen model of recombination hotspots evolution in the light of archaic and modern human genomes. PLoS genetics 10(11):e1004790.

38. Fledel-Alon A, et al. (2011) Variation in human recombination rates and its genetic determinants. PloS one 6(6):e20321.

39. Myers S, Bottolo L, Freeman C, McVean G, & Donnelly P (2005) A finescale map of recombination rates and hotspots across the human genome. Science 310(5746):321–324.

40. Sankararaman S, et al. (2014) The genomic landscape of Neanderthal ancestry in present-day humans. Nature 507(7492):354–357.

41. Vernot B & Akey JM (2014) Resurrecting surviving Neandertal lineages from modern human genomes. Science 343(6174):1017–1021.

42. McVicker G, Gordon D, Davis C, & Green P (2009) Widespread genomic signatures of natural selection in hominid evolution. PLoS genetics 5(5):e1000471.

43. Charlesworth B, Morgan M, & Charlesworth D (1993) The effect of deleterious mutations on neutral molecular variation. Genetics 134(4):1289–1303.

44. Cai JJ, Macpherson JM, Sella G, & Petrov DA (2009) Pervasive hitchhiking at coding and regulatory sites in humans. PLoS genetics 5(1):e1000336.

45. Busing FM, Meijer E, & Van Der Leeden R (1999) Delete-m jackknife for unequal m. Statistics and Computing 9(1):3–8.

46. Wood R (2015) From revolution to convention: the past, present and future of radiocarbon dating. Journal of Archaeological Science 56:61–72.

47. Reich D, et al. (2011) Denisova admixture and the first modern human dispersals into Southeast Asia and Oceania. The American Journal of Human Genetics 89(4):516–528.

48. Meyer M, et al. (2012) A high-coverage genome sequence from an archaic Denisovan individual. Science 338(6104):222–226.

49. Lipson M, et al. (2015) Calibrating the Human Mutation Rate via Ancestral Recombination Density in Diploid Genomes. bioRxiv:015560.

50. Coop G, Wen X, Ober C, Pritchard JK, & Przeworski M (2008) Highresolution mapping of crossovers reveals extensive variation in fine-scale recombination patterns among humans. Science 319(5868):1395–1398.

51. Fenner JN (2005) Cross-cultural estimation of the human generation interval for use in genetics based population divergence studies. American journal of physical anthropology 128(2):415–423.

52. Helgason A, Hrafnkelsson B, Gulcher JR, Ward R, & Stefánsson K (2003) A populationwide coalescent analysis of Icelandic matrilineal and patrilineal genealogies: evidence for a faster evolutionary rate of mtDNA lineages than Y chromosomes. The American Journal of Human Genetics 72(6):1370–1388.

53. Hudson RR (2002) Generating samples under a Wright–Fisher neutral model of genetic variation. Bioinformatics 18(2):337–338.

54. Patterson N, Price AL, & Reich D (2006) Population structure and eigenanalysis. PLoS genetics 2(12):e190.

55. Patterson N, et al. (2012) Ancient Admixture in Human History. Genetics 192(3):1065–1093.

